# Draft genome of the liver fluke *Fasciola gigantica*

**DOI:** 10.1101/451476

**Authors:** Tripti Tripathi, Arpita Ghosh, Vivek N Todur, Parismita Kalita, R Vijayakumar, Jupitara Kalita, Rohit Shukla, Purna B Chetri, Harish Shukla, Amit Sonkar, Denzelle Lee Lyngdoh, Radhika Singh, Surendra K Chikara, Timir Tripathi

## Abstract

Fascioliasis is a neglected food-borne disease caused by liver flukes (genus *Fasciola*) and affects more than 200 million people worldwide. Despite technological advances, little is known about the molecular biology and biochemistry of the fluke. We present the draft genome of *Fasciola gigantica* for the first time. The assembled draft genome has a size of ~1.04 Gb with an N50 of 129 kb. A total of 20,858 genes were predicted. The *de novo* repeats identified in the draft genome were 46.85%. In pathway analysis, all the genes of glycolysis, Kreb’s cycle and fatty acid metabolism were found to be present, but the key genes for fatty acid production in fatty acid biosynthesis were missing. This indicates that the fatty acid required for the survival of the fluke may be acquired from the host bile. The genomic information will provide a comprehensive resource to facilitate the development of novel interventions for fascioliasis control.

## INTRODUCTION

Fascioliasis, caused by the trematodes of the genus *Fasciola*, is an important food-borne parasitic disease belonging to the group of neglected tropical diseases (NTDs). Food-borne trematodiases are one of the 17 NTDs included in the WHO roadmap for lowering the public-health burden of NTDs ^1^. *Fasciola hepatica* and/or *Fasciola gigantica* infection is prevalent in over 600 million domestic ruminants (cattle, sheep, pig, donkey, buffalo, and goats), causing major economic losses previously estimated to be US$3 billion p.a.^2^. Fascioliasis has remarkable latitudinal, longitudinal and altitudinal distribution given that these liver flukes can adapt to different environments and habitats including extreme climatic conditions. *F. gigantica* is found in the tropical regions of Africa, Asia and the Middle East, where it affects 25–100% of total cattle populations. It is also prevalent in the livestock populations of India, Pakistan, Indonesia, Indochina, and the Philippines. Human infection has been reported in 51 different countries from five continents; this indicates the geographical expansion of the problem ^3-6^. In humans, fascioliasis is an important zoonotic disease that has been estimated to infect 2.4–17 million people and has put approximately 180 million people at risk globally ^7-9^. Major human fascioliasis endemic areas include Africa, Europe, the Middle East (including Egypt), Southeast Asia, and Latin America, with the highest prevalence at 72–100% in the Bolivian Altiplano ^10,11^. Interestingly, it has been observed that in hyper-endemic areas, the parasite is better adapted to the human host^3^. Most cases of human fascioliasis are reported on *F. hepatica* ^3,6,10,11^, though a few reports on *F. gigantica* causing human infection are available^12-14^.

The adult *F. gigantica* is hermaphroditic containing both male and female sexual organs, and capable of self-fertilization. The life cycle of *Fasciola* involves an intermediate host—snail of the family Lymnaeidae—and a mammalian definitive host. Infection starts on ingesting food contaminated with the larval stage, i.e., metacercariae, which are found floating freely in fresh water or attached to water plants. The metacercariae excyst in the duodenum of the mammalian host and then migrate to the liver through the intestinal wall; the adults mature in the biliary ducts. The eggs are passed into the intestine and then excreted out through feces ^3^. Clinical symptoms of fascioliasis are caused by the migration of the young flukes through the liver causing abdominal pain, weight loss, fever, nausea, vomiting, hepatomegaly, hepatic tenderness, and eosinophilia. The infection causes extensive damage to the liver and may lead to portal cirrhosis. Long-term infection by *Fasciola* results in chronic stimulation of the bile duct epithelium due to the excretory-secretory (ES) products released from parasites into the host bile environment ^15^. These ES products have key roles in feeding behavior, detoxification of bile components, and immune evasion by liver flukes ^15^. Transcriptome data sets for *F. gigantica* include substantial representation of ES products, suggesting its role in the mechanism of infection of this parasite ^16^. The WHO has recommended triclabendazole, a benzimidazole compound, as the drug of choice for the treatment of fascioliasis as it is active against key parasite stages, i.e. early juvenile, juvenile and adult stages. However, recent studies have suggested that the *F. hepatica* have gained resistance to triclabendazole in several countries^17-20^. In principle, food-borne trematodes can be effectively controlled using multiple interventions implemented simultaneously across sectors.

Recently genomes from the parasitic flukes including *Schistosoma japonicum ^21^, S. mansoni*^22^, *S. haematobium*^23,24^, *Opisthorchis viverrni*^25^, *Clonorchis sinensis*^26,27^ and *F. hepatica*^28,29^ have been sequenced. The genome sequences shed light on how these organisms survive in the host environment and show that the metabolic pathways in the parasite are highly adapted to the host conditions. Here, we report the draft sequence, assembly, and analysis of the *F. gigantica* genome. It is one of the largest parasitic genomes to be sequenced. The genomic information provides a resource to facilitate development of novel interventions for fascioliasis control.

## RESULTS AND DISCUSSION

### *De novo* genome assembly and annotation

To avoid technical difficulties in assembly, genomic DNA was isolated from a single adult fluke, and one each of shotgun sequencing library and Mate-pair DNA library were constructed with a library size of approximately 350 bp. The Paired-end and Mate-pair libraries were sequenced using HiSeq 2500 to generate 32.7 Gb and 1.7 Gb of data, respectively. The raw reads were quality filtered and adapter trimmed. The filtered high-quality reads were assembled using SOAPdenovo (v1.5.2) program. This primary assembly was further used for gap filling by Paired-end and Mate-pair reads using GapCloser. Further, SSPACE v2.0 was used for scaffolding. The resultant assembly was used in Chromosomer v0.1.4a for further improvement of the assembly. The assembled draft genome obtained was 40,381 scaffolds with a genome size of 1.04 Gb (Table 1). The genome size was similar to *F. hepatica* genome and much larger than the genomes of other parasitic flukes (Table 2). A total of 16,465 scaffolds were larger than 10 Kb size, resulting in 978.97 Mb of genome length comprising of 94.11% of the genome assembly. The completeness of the genome was estimated to be 51.3 % and fragements is estimated to be 12.3% using BUSCO2.0. The BUSCO completeness of the published genomes of Platyhelminthes was found to be ranging from 20% to 73% as reported in WormBase database (http://parasite.wormbase.org/species.html#Platyhelminthes).

**Table 1.**
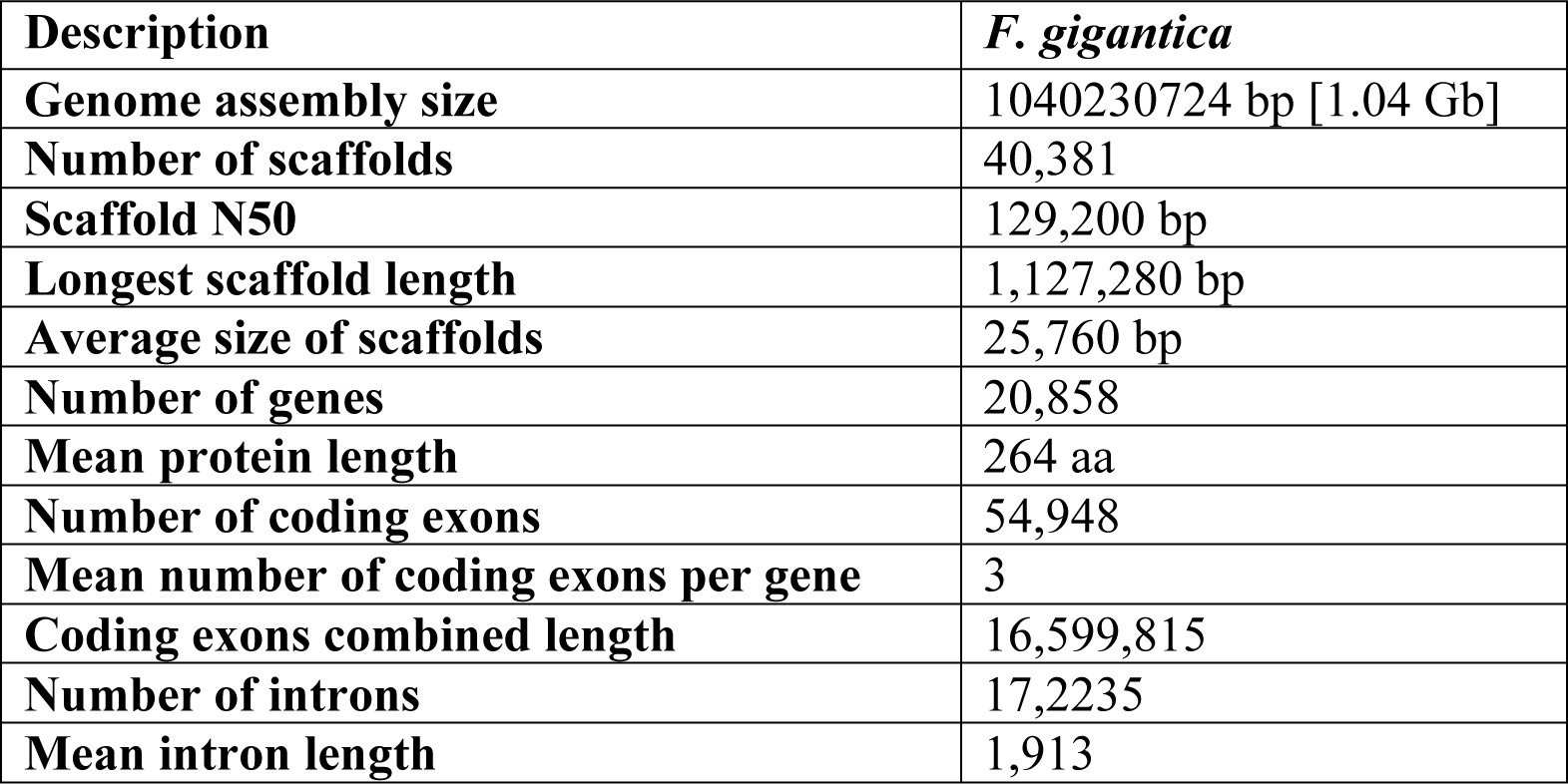
The assembly features.

**Table 2.** Comparison of the nuclear genome assemblies of *F. gigantica* and other parasitic flukes.

|  | *F. gigantica*<br>(Present work) | *F. hepatica*<sup>29</sup> | *F. hepatica*<sup>28</sup> | *O. viverrini*<sup>25</sup> | *C. sinensis*<sup>26,27</sup> | *S. japonicum*<sup>21</sup> | *S. haematobium*<sup>23,24</sup> | *S. mansoni*<sup>22</sup> |
| --- | --- | --- | --- | --- | --- | --- | --- | --- |
| **Genome size** | 1.04 Gb | 1.13 Gb | 1.27 Gb | 634.5 Mb | 320.5 Mb | 397 Mb | 385 Mb | 364 Mb |
| **Number of genes** | 20858 | 14,851 | 22,676 | 16379 | 28407 | 13469 | 13073 | 13184 |
| **Mean number of exons per gene** | 3 | 3.18 | 5.3 | 5.8 | 7.7 | 5.3 | 5.4 | 6 |
| **Mean exon length (bp)** | 3025 | 257 | 303 | 254 | 312 | 222 | 246 | 222 |
| **Mean intron length (bp)** | 1913 | NA | 3700 | 3531 | 359 | 2059 | 2442 | 2407 |
| **Total GC content (%)** | 43.76 | 41 | 44 | 47.8 | 34.06 | 33.5 | 34.3 | 34.7 |

### Repeat annotation

The repeats were detected using denovo method. The *de novo* predicted *F. gigantica* specific repeats were 487,374,279 bp, accounting for 46.85% of the entire genome. The total numbers of repeat sequences identified were represented in 40,381 scaffolds. The repeat unit length ranged from 12 to 2,253,045 bp. We have identified 21.26% LINEs, 6.76% LTR elements, 45.93% total interspersed repeats and 15.09% of unclassified repeats, as summarized in Table 3. The details of the repeats are provided in Supplementary Table S1.

**Table 3.** Summary of the *de novo* repeats identified.

| **Description** | **Number of elements** | **Length occupied in bp** |
| --- | --- | --- |
| **SINEs:** | 67024 | 11130748 bp |
| **MIRs** | 1689 | 186234 bp |
| **LINEs:** | 469965 | 221172212 bp |
| **LINE2** | 12439 | 4408849 bp |
| **L3/CR1** | 134130 | 66193633 bp |
| **LTR elements** | 130496 | 70357405 bp |
| **ERV_classI** | 344 | 32937 bp |
| **ERV_classII** | 2413 | 579624 bp |
| **DNA elements** | 63140 | 18072908 bp |
| **TcMar-Tigger** | 176 | 53598 bp |
| **Unclassified** | 697673 | 157005240 bp |
| **Total interspersed repeats** |  | 477738513 bp |
| **Small RNA** | 35107 | 6310113 bp |
| **Satellites** | 45457 | 7519158 bp |
| **Simple Repeats** | 68849 | 3254749 bp |
| **Low Complexity** | 2024 | 92148 bp |

### Gene prediction and annotation

This draft genome was further used for gene prediction using *Schistosoma* as model species. The gene prediction method was used to identify the protein coding genes. A total of 20,858 genes were predicted with an average gene length of 795 bp and 264 aa.Of them, 59% (12,285 genes) were found to have homology with NCBI NR database and 13.9% (2,900 genes) were classified with GO terms (details provided in Supplementary Table S2). The annotation of genes showed the highest hits against *F. hepatica* (5,248), followed by *O. viverrini* (1,389). A total of 5,371 genes were annotated according to all the three GO sub vocabularies (i.e. cellular component (CC), biological process (BP), and molecular function (MF)). A total of 2,013 genes were classified in BP; 2,352 genes as MF; and 1,276 genes as CC. Out of total 20,858 genes, 807 genes have been found to have all three categories of GO terms (Figures 1 and 2). Genes associated with similar functions were assigned to the same GO functional group. Further, the proteins for *F. gigantica* and *F. hepatica* were compared using Blast with 90% identity. The protein coding genes in these two genomes were found to have 65.3% similarity. Out of the total genes which were similar in both the genome, only 3688 genes were found to have GO terms, which included 1403 cellular component (CC), 2474 biological process (BP) and 3143 molecular function (MF), the details are mentioned in Supplementary Figure S1.

**Figure 1.**
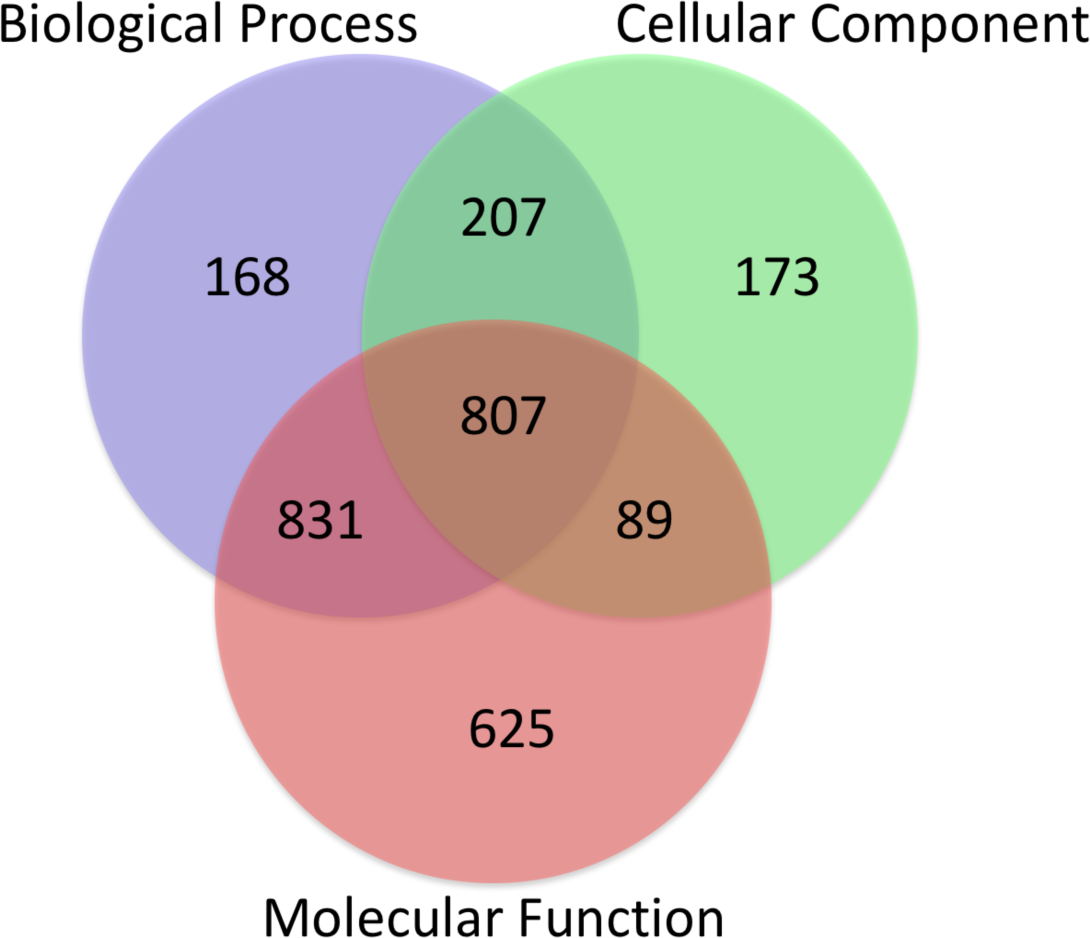
Graphical representation of distribution of genes assigned to GO terms. Proportion of the 5,371 *F. gigantica* proteins with functional information in different GO categories is shown as, Biological Process, Molecular Function and Cellular Component.

**Figure 2.**
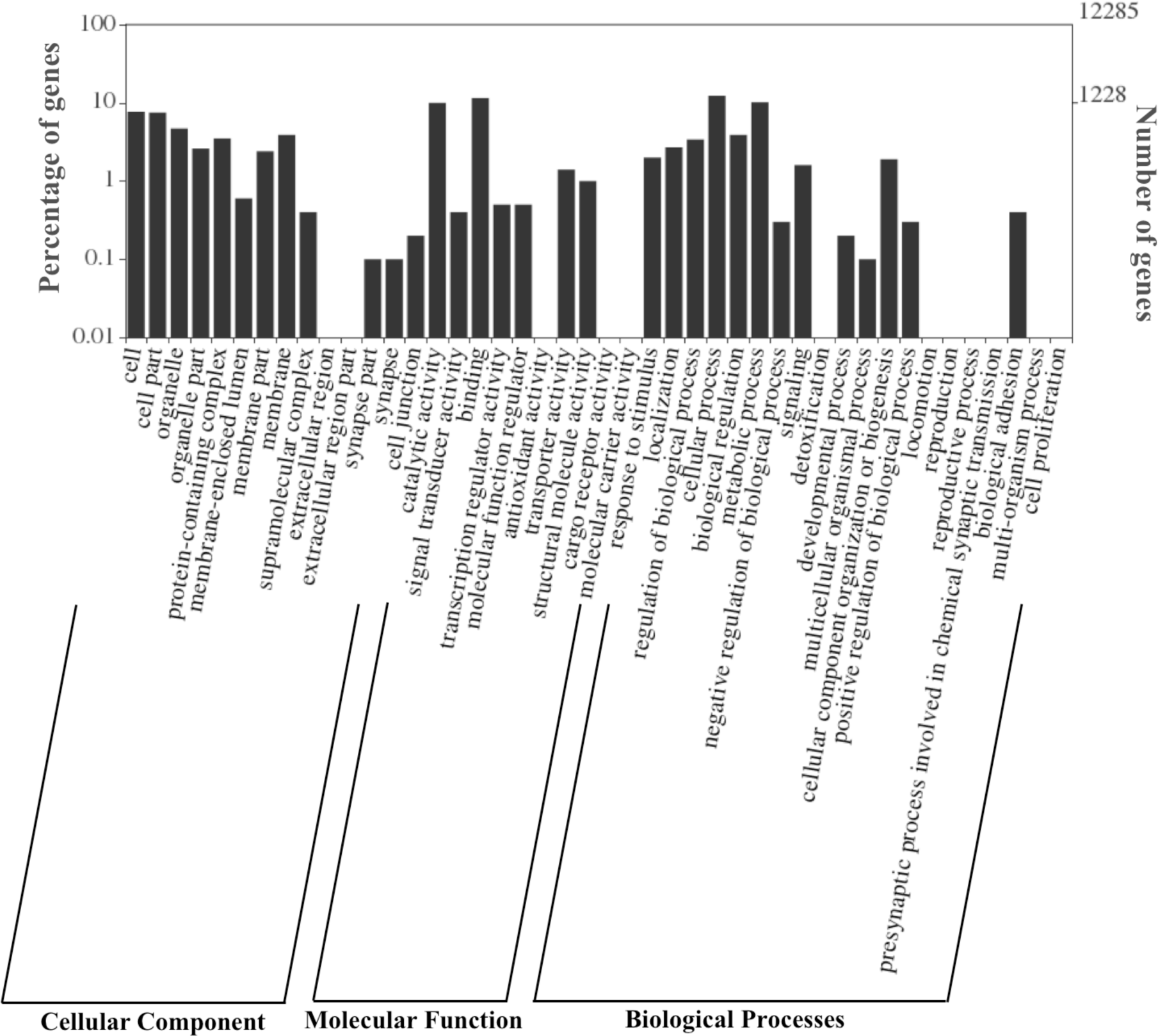
GO classification of genes in cellular components, molecular function and biological process.

The ES proteins found were cathepsin proteases (which includes cathepsin L-like proteases, cathepsin B-like proteases and cathepsin D-like proteases), glutathione transferase, fatty acid-binding protein and glyceraldehyde-3-phosphate dehydrogenase ^15^. A total of 23 blast hits against ES protiens were identified from the Blast results, in which predominantly, cathepsin protein was found (Supplementary Table S2 : sheet NR annotation). Cathepsin B and L cysteine proteases are known to be important antigens produced in the trematode mainly in genus *Fasciola* and play an important role in parasite nutrition, immune evasion and host invasion^30^. Total of 46 GO terms were assigned and 4 genes had missing GO terms (Supplementary Table S2). The proteins which were significantly enriched are classified in the following GO terms, proteolysis, cysteine-type endopeptidase activity and regulation of catalytic activity 29. The GO terms of ES proteins were classified in 0 CC, 19 MP and 13 BP. Earlier studies have suggested that the cathepsins help the parasite to survive inside the host gall bladder and bile duct. Trematodes encode various subfamilies of cathepsins, which in turn provide insight of the host parasite relationships and developmentally regulated expression with the passage of the parasites through the host in the life cycle ^31^. Proteases may help in the activation of cathepsins which in turn facilitate the digestion of host tissues, releasing essential amino acids^22^.

### Annotation of conserved domains

The search made against InterPro database provided 14,487 InterPro hits, 4,810 InterPro hits with GO terms and 6,371 no hits. The GO terms in Interpro were merged, which resulted in GO before merge 9,039, GO after merge 12,285, confirmed IPS GO 20,351 and too general IPS GO 1,608.

The analysis revealed that 5,205 protein sequences were categorized into 1,591 domains and 2,448 families. InterPro domains/families were sorted according to the gene sequences assigned; distribution of the top 20 InterPro domains has been represented in Figure 3. The most abundant domain (IPR000477), i.e., reverse transcriptase domain, was obtained with 1,155 annotated gene sequences, followed by (IPR001584) Integrase, catalytic core with 235 annotated gene sequences and (IPR000719) Protein kinase domain with 481 annotated gene sequences. The InterPro families distribution are represented in Figure 4, and the top 5 families identified are (IPR036691) Endonuclease/exonuclease/phosphatase superfamily, (IPR027124) SWR1-complex protein 5/Craniofacial development protein, (IPR027417) P-loop containing nucleoside triphosphate hydrolase, (IPR036397) Ribonuclease H superfamily and (IPR012337) Ribonuclease H-like superfamily.

**Figure 3.**
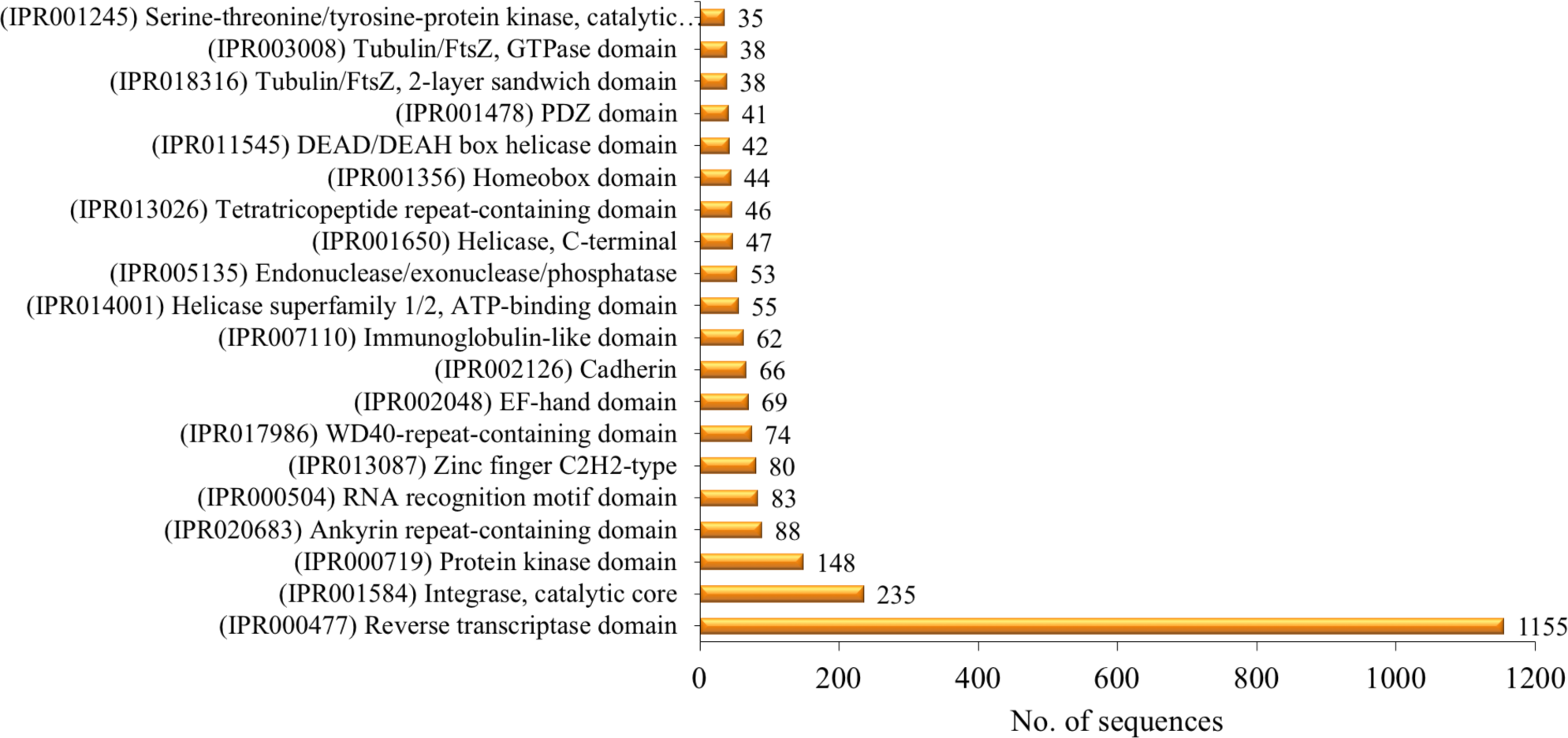
The 20 most abundant InterPro domains revealed by InterProScan annotation are represented. The most abundant domain (IPR000477) Reverse transcriptase domain was obtained with 1,056 annotated gene sequences.

**Figure 4.**
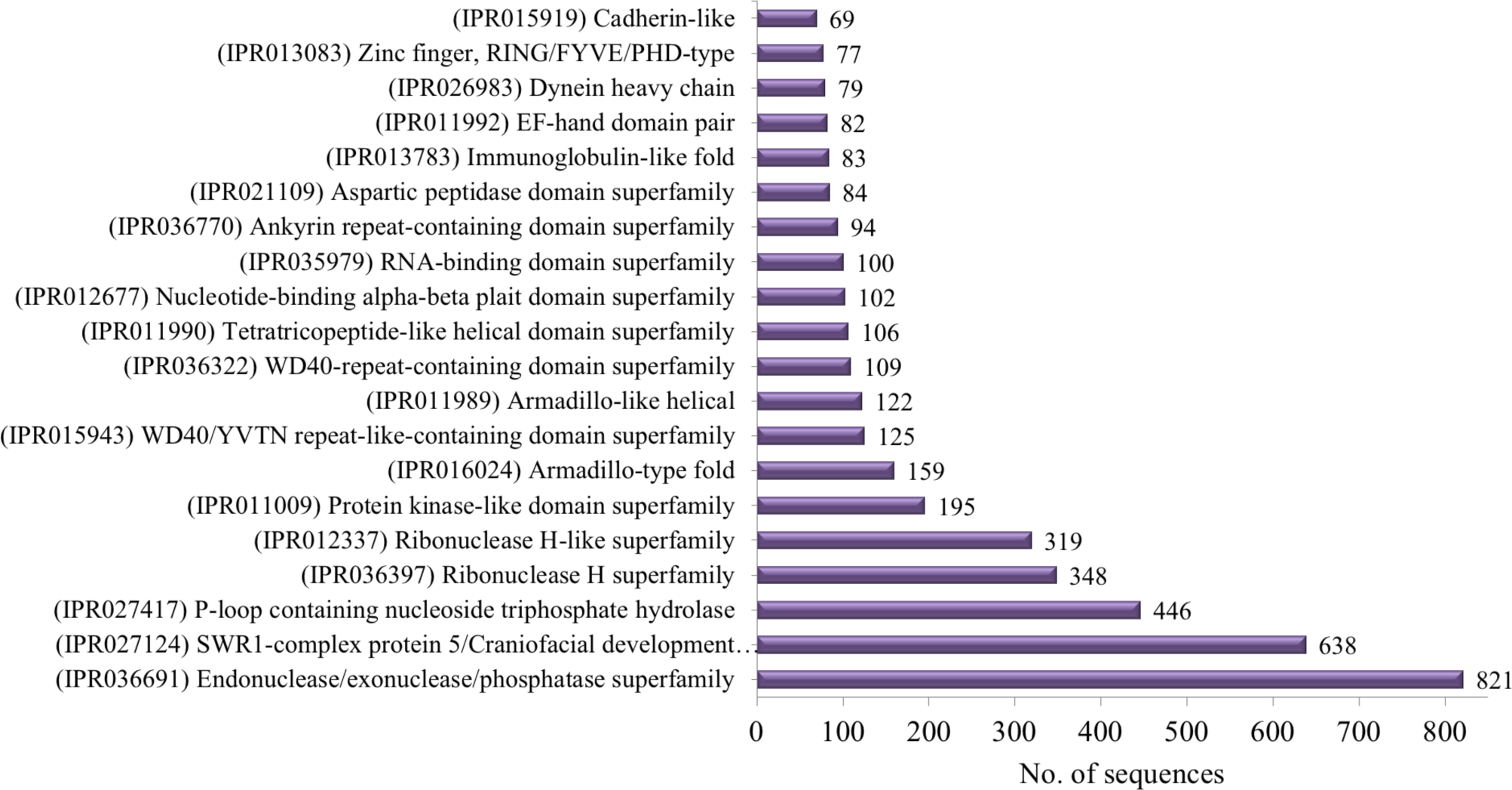
The 20 most abundant InterPro families revealed by InterProScan annotation are represented. The most abundant family identified was (IPR027124) SWR1-complex protein 5/Craniofacial development protein.

The search found 2,084 Pfam domains in 6,693 genes, in which the reverse transcriptase domain [PF00078] and Integrase, catalytic core [PF00665] domains were highly represented by 989 and 175 genes, respectively. The details of the conserved domains/families are provided in Supplementary Table S3.

### Pathway analysis

KAAS was used to carry out ortholog and mapping of the genes to the biological pathways. The annotated genes were compared against those available in the KEGG database using BLASTx with default threshold bit score value and expected threshold. Total assigned KO ID was 1,343 of 4,016 genes that were mapped to respective pathways (details provided in Supplementary Table S2). The mapped genes represented a metabolic pathway of major biomolecules such as carbohydrates, amino acids and other pathways.

According to the life cycle of *F. gigantica*, they can obtain energy from both aerobic and anaerobic metabolism ^32^. The adult metabolism is anaerobic, and juvenile metabolism is almost aerobic. From the life cycle, it is evident that all liver flukes inhabit the bile duct, which is anaerobic, but for the survival in the intermediate host, biochemical pathways of aerobic metabolism play crucial roles. The glycolytic pathway shows the presence of all the key enzymes, such as hexokinase [EC:2.7.1.1], enolase [EC:4.2.1.11], pyruvate kinase [EC:2.7.1.40], and lactate dehydrogenase [EC:1.1.1.27] (Supplementary Fig. S2). Some of the genes involved in energy metabolism were absent, indicating that the adult worms utilize the glucose exogenously from the glycolytic pathway or may absorb nutrients from the host under anaerobic conditions ^26^. The Kreb’s cycle is also an important pathway for energy metabolism (Supplementary Fig. S3).

In fatty acid metabolism pathway, all the genes encoding enzymes are present (Supplementary Fig. S4). In contrast, for the fatty acid biosynthesis pathway, only three enzymes-acetyl-CoA carboxylase / biotin carboxylase 1 [EC:6.4.1.2 6.3.4.14], 3-oxoacyl-[acyl-carrier-protein] synthase II [EC:2.3.1.179] and long-chain acyl-CoA synthetase [EC:6.2.1.3] were present (Supplementary Fig. S5). It is known that the fatty acid binding proteins in liver flukes play a crucial role in utilizing the fatty acid produced by the host bile. Therefore, liver flukes do not need to synthesize their own fatty acids endogenously^26^. The key genes are present in fatty acid metabolism but are absent in the fatty acid biosynthesis pathway, which reflects that liver fluke receives fatty acids from the host bile.

### Analysis of orthologous groups

*F. gigantica* and *F. hepatica* genomes were predicted to have 20,858 and 33,454 proteins, which resulted in 9,365 clusters. A total of 6,241 core genes (i.e., in the cluster multiple copies of genes are present) and 5,654 single copies of gene clusters were identified between the two genomes using OrthoVenn (Figure 5A). Unique ortholog clusters 905 and 2,219 were deciphered in *F. gigantica* and *F. hepatica* genome, respectively.

**Figure 5.**
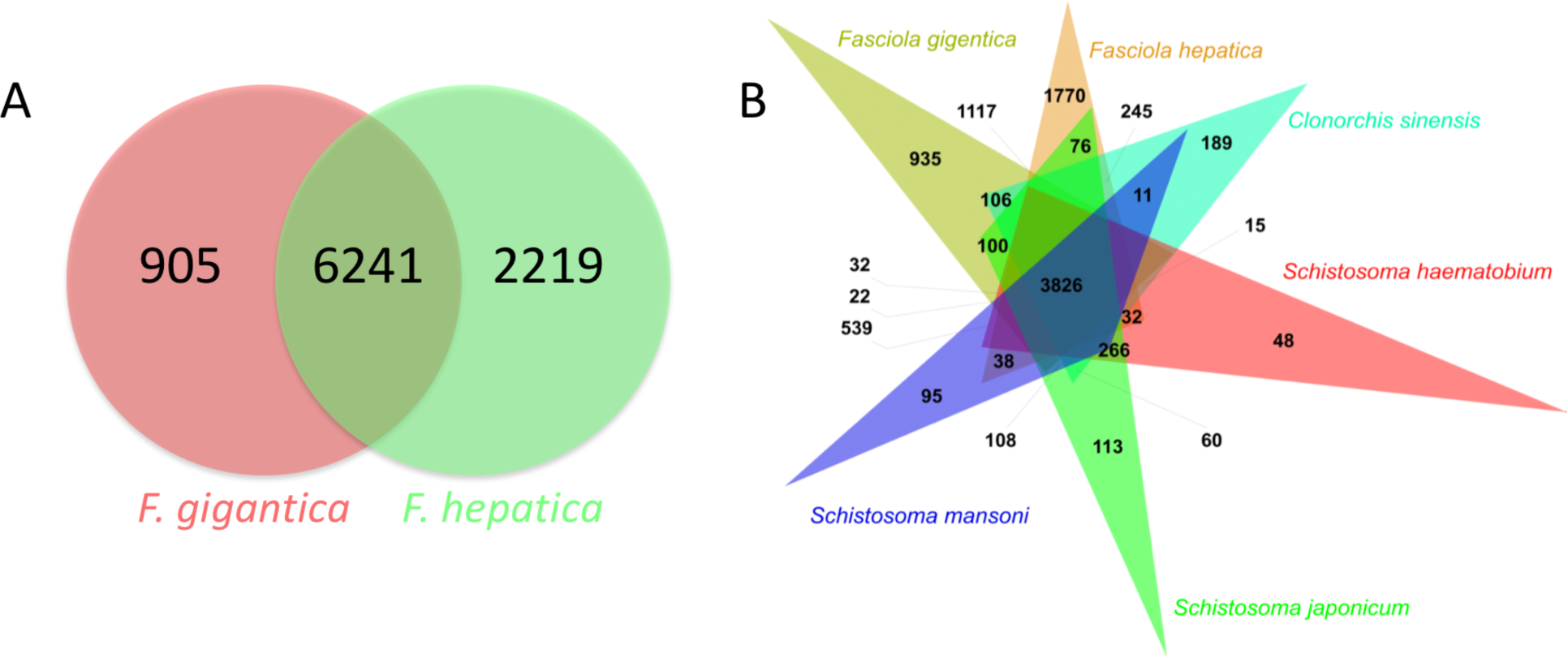
Venn diagram showing phylogenetic distribution of orthologous protein families. ***A.*** Between *F. gigantica* and *F. hepatica*. ***B.*** Between *F. hepatica, F. gigantica, S. mansoni* and *O. viverrini*.

Similarly, we compared six genomes, i.e., *F. gigantica, F. hepatica, S. mansoni, S. japonicum, S. haematobium and C. sinensis*. The total predicted proteins for S. mansoni, S. japonicum, S. haematobium, and C. sinensis were 11774, 12738, 11140 and 13634, respectively. Total clusters generated were 14,288 out of which 11,138 orthologous clusters were common in at least two species and 1,863 single copies genes clusters. The total number of clusters identified in each genome is 7,664, 10,289, 8,298, 8,010, 8,455 and 7,664 respectively. The core genes identified were 3,826 from all the six species as shown in Figure 5B. The unique orthologous clusters identified in *F. gigantica, F. hepatica, S. mansoni, S. japonicum, S. haematobium and C. sinensis* were 935, 1770, 95, 113, 48 and 189, respectively. The details are provided in Supplementary Table S4.

## CONCLUSIONS

The liver fluke *F. gigantica* is a major parasite of livestock worldwide, causing huge economic losses to agriculture, as well as 2.4-17 million human infections annually. We studied the draft genome of the organism, which is among the largest known parasitic genomes at 1.3 Gb. The larger genome size may be attributed to gene duplication and polymorphism that may help adaptation of the parasite to the host environment and the capacity for rapid evolution. The genomic information will provide new insights into its adaptation to the host environment, external selection pressures and will help in the development of novel therapies for fascioliasis control.

## METHODS

### DNA isolation

*F. gigantica* flukes were collected from the liver of naturally infected cattle from the Bara Bazar slaughter house, Shillong, India (Latitude- 25.5724472; longitude- 91.8745219). The whole worm was washed with 70% ethanol followed by rinsing several times with 1x phosphate buffer saline. Individual flukes were immediately frozen in liquid nitrogen and stored at −80°C until processed for genomic DNA extraction. A single individual worm was crushed in liquid nitrogen to isolate its genomic DNA using the standard phenol–chloroform extraction method. Quality and integrity of the isolated DNA were checked on 0.8% Agarose gel and Nano-drop spectrofluorimeter.

### DNA library construction and sequencing

One shotgun sequencing library and one Mate-pair DNA libraries were constructed according to the Illumina Sample Preparation Guide (Illumina, San Diego, CA, USA). The shotgun paired-End sequencing library with insert size of approximately 350 bp was prepared using TruSeq Nano DNA Library Prep Kit for Illumina. Briefly, 200 ng of DNA was fragmented by Covaris M220 to generate a mean fragment distribution of 300-400 bp. Covaris shearing generates dsDNA fragments with 3′ or 5′ overhangs that were then subjected to End Repair Mix to convert the overhangs into blunt ends. The 3′ to 5′ exonuclease activity of this mix removes the 3′ overhangs, and the 5′ to 3′ polymerase activity fills in the 5′ overhangs. A single ‘A’ base was then added to the ends of the polished DNA fragments followed by adapter ligation to ensure a low formation rate of chimera (concatenated template). Indexing adapters were ligated to the ends of the DNA fragments to prepare them for hybridization onto a flow cell. The ligated products were size selected using Agencourt AMPure XP beads (Beckman Coulter Life Sciences, USA) and PCR enriched with the Illumina adaptor index PCR primer for six cycles.

The Mate-pair sequencing library was prepared using Illumina Nextera Mate-pair Sample Preparation Kit. Briefly, 4 µg of the high-quality gDNA was tagmented using Mate-pair transposomes. Using Zymo Genomic DNA Clean & Concentrator kit (Zymo Research, USA) the tagmented DNA was purified and then fragmented for circularization by repairing the ends by strand displacement reactions. Short fragments less than 1500 bp were removed using Ampure XP bead clean up steps. Precise size selection was carried out using Pippin prep system to select 8-11 kb fragments followed by clean up using Zymo clean Genomic DNA Clean & Concentrator Kit. The DNA fragments were then self-circularized by an intramolecular ligation, and non-circularized DNA was removed by DNA exonuclease treatment. The large circularized DNA fragments were physically sheared to smaller sized fragments (approximately 300–1000 bp) in Covaris using defined shearing parameter. The sheared DNA fragments (Mate-pair fragments) containing the biotinylated junction adapter were purified by binding to streptavidin magnetic beads, and the unwanted, un-biotinylated molecules were removed through a series of washes. The streptavidin bead bound fragments were then subjected to end repair, A-tailing, Illumina adapter ligation, and final PCR enrichment for the mate pair fragments that have TruSeq DNA adaptors on both the ends.

The library validation was carried out using Tape Station 4200 (Agilent Technologies) using D1000 Screen Tape assay kit. The Paired-end sequencing run was performed on HiSeq 2500 (Illumina) using 2X125 bp read chemistry.

### Genome assembly

The whole genome sequencing was carried out for Paired-end and Mate-pair library using HiSeq 2500 with 2×125 bp chemistry. The raw Mate-pair reads were extracted using in-house script based on their orientation and presence of junction adapter in between read1 and read2. The reads having junction adapter in between the reads were used as Mate-pair reads ^33^. The raw reads were adapter trimmed and quality filtered using Trimmomatic (v 0.35) ^34^ with a minimum read length cut-off of 100 bp. The assembly of Paired-end and Mate-pair reads was carried out using SOAPDenovo (v1.5.2) with optimized 57 kmer length. After the primary assembly, GapCloser was used for gap filling and scaffolding with both Paired-end and Mate-pair libraries. Further, scaffolding was carried out using SSPACE v2.0^5^. The resultant assembly was used with the available (ftp://ftp.ncbi.nlm.nih.gov/genomes/all/GCA/002/763/495/GCA_002763495.1_F_hepatica_1.0.allpaths.pg) using Chromosomer v0.1.4a^5^. The assembled draft genome was used in downstream analysis. The completeness of the genome was estimated using BUSCO2.0. *De novo* repeat identification was performed using RepeatModeler. The *de novo* repeat libraries were constructed using the draft genome with RepeatModeler, which contains two repeat finding programs (RECON and RepatScout). This resulted in a repeat library with classified repeat families that was used in RepeatMasker as repeat library, on the draft genome to identify the *de novo* repeats.

### Gene annotation

The draft genome of *F. gigantica* was used for gene prediction using Augustus v3.2.1 ^35^ with the gene model parameters tuned for *Schistosoma*; the rest parameters were kept as default. Functional annotations of the predicted genes were performed using BLASTx program keeping an e value 1e-6 against the NCBI NR database. BLASTx finds the homologous sequences for the genes against NR database. Homologs of *F. gigantica*-predicted protein sequences were identified using BLAST, and the functional domains were identified using InterPro. The results of BLAST searches were used as an input to Blast2GO PRO ^36^. Based on the BLAST hits obtained, gene ontology (GO) annotation was performed to obtain the GO terms and classify them into biological process, molecular functions and cellular components. The GO terms associated with each of the BLAST results (mapping step) and the GO annotation assigned to the query (annotation step) were obtained. Further, the conserved domain/motifs were identified using InterProScan (IPS), an online plugin of BLAST2GO that combines various protein signature recognition methods with the Interpro database. The resulting GO terms were merged with the GO term results obtained from the above annotation step. The protein coding gene sequences of *F. gigantica* and *F. hepatica* (PRJEB6687) (downloaded from WormBase WBPS10 :http://parasite.wormbase.org) were aligned using Blastn to identify the similarity in the protein coding genes. The *F. hepatica* genes were used as a database for the Blast against *F. gigantica* protein with an e value of 1e-5.

### Pathway analysis

To identify the potential involvement of the predicted genes of *F. gigantica* in biological pathways, the predicted genes were aligned to KEGG pathway database using KEGG (Kyoto Encyclopaedia of Genes and Genomes) automatic annotation server ^37-39^. KEGG analysis includes KEGG Orthology (KO) assignments and Corresponding Enzyme commission (EC) numbers and metabolic pathways of predicted genes using KEGG automated annotation server KAAS (http://www.genome.jp/kaas-bin/kaas_main). The genes distribution under the respective EC number was used to map them to the KEGG biochemical pathways. This process provides an overview of the different metabolic processes active within an organism and enables further understanding of the biological functions of the genes.

### Identification of orthologous groups

The protein sequences of *F. hepatica, S. mansoni, S. japonicum, S. haematobium and C. sinensis*were obtained from WormBase Parasite database (http://parasite.wormbase.org). Protein sequences of *F. gigantica* and *F. hepatica* were used to perform an all against all comparison using BLASTP with orthoVenn at default parameters ^40^. The core genes and unique genes were identified between *F. gigantica* and *F. hepatica* genomes. The ortholog analysis was also performed with *F. hepatica, S. mansoni, S. japonicum, S. haematobium and C. sinensis*. This enabled us to elucidate the function and evolution of protein across the six species.

## Accession numbers

This Whole Genome Shotgun project has been deposited at DDBJ/ENA/GenBank under the accession MKHB00000000. The version described in this paper is version MKHB02000000.

## Authors’ contributions

Conceived and designed the experiments: TrT, AG, TiT.

Performed the experiments: TrT, AG.

Analyzed the data: TrT, AG, VT, SKC, TiT.

Contributed reagents/materials/analysis tools: TrT, AG, SKC, PK, VR, JK, PBC, RoS, DL, HS, AS, RaS.

Wrote the paper: TiT, AG.

